# A combination of phenotypic responses and genetic adaptations enables *Staphylococcus aureus* to withstand inhibitory molecules secreted by *Pseudomonas aeruginosa*

**DOI:** 10.1101/2025.02.24.639796

**Authors:** Lukas Schwyter, Selina Niggli, Natalia Zajac, Lucy Poveda, Witold Eryk Wolski, Christian Panse, Ralph Schlapbach, Rolf Kümmerli, Jonas Grossmann

## Abstract

*Staphylococcus aureus* and *Pseudomonas aeruginosa* frequently co-occur in infections, and there is evidence that their interactions can negatively affect disease outcomes. *P. aeruginosa* is known to be dominant, often compromising *S. aureus* through the secretion of inhibitory compounds. We previously demonstrated that *S. aureus* can become resistant to growth-inhibitory compounds during experimental evolution. While resistance arose rapidly, the underlying mechanisms were not obvious as only a few genetic mutations were associated with resistance, while ample phenotypic changes occurred. We thus hypothesize that resistance may result from a combination of phenotypic responses and genetic adaptation. Here, we tested this hypothesis using proteomics. We first focused on an evolved strain that acquired a single mutation in *tcyA* (encoding a transmembrane transporter unit) upon exposure to *P. aeruginosa* supernatant. We show that this mutation leads to a complete abolishment of transporter synthesis, which confers moderate protection against PQS and selenocystine, two toxic compounds produced by *P. aeruginosa*. However, this genetic effect was minor compared to the fundamental phenotypic changes observed at the proteome level when both ancestral and evolved *S. aureus* strains were exposed to *P. aeruginosa* supernatant. Major changes involved the downregulation of virulence factors, metabolic pathways, membrane transporters, and the upregulation of ROS scavengers and an efflux pump. Our results suggest that the observed multi-variate phenotypic response is a powerful adaptive strategy, offering instant protection against competitors in fluctuating environments and reducing the need for hard-wired genetic adaptations.

**Importance:** Different bacterial pathogens can co-occur in infections, where they interact with one another and influence disease severity. Previous research showed that pathogens can evolve and adapt to co-infecting species. Here, we show that evolution through genetic mutations and selection are not necessarily required to change pathogen behavior. Instead, we found that the human pathogen *Staphylococcus aureus* is able to plastically respond to the presence of *Pseudomonas aeruginosa*, a competing pathogen. Through proteomics and metabolomics, we demonstrate that *S. aureus* undergoes substantial proteomic alterations in response to *P. aeruginosa* by down-regulating virulence factor expression, changing metabolism, and mounting protective measures against toxic compounds. Our work highlights that pathogens possess sophisticated mechanisms to respond to competitors to secure growth and survival in polymicrobial infections. We predict such plastic responses to have significant impacts on infection outcomes.

## Introduction

Polymicrobial infections involving *Pseudomonas aeruginosa* and *Staphylococcus aureus* are frequently observed in clinical settings, especially in respiratory, wound, and bloodstream infections (1–4). These co-infections often result in more severe disease outcomes than mono-species infections (5, 6). Notably, *P. aeruginosa* and *S. aureus* are commonly found together in the lungs of cystic fibrosis (CF) patients, where their interactions are associated with increased lung damage and chronic infection (7, 8). The interplay between *P. aeruginosa* and *S. aureus* is complex and can significantly influence infection dynamics and virulence, and complicate treatment strategies (9, 10). Co-infections with *P. aeruginosa* and *S. aureus* can lead to increased antibiotic resistance, often necessitating more aggressive or combined treatment approaches (11).

Laboratory studies typically find *P. aeruginosa* to be dominant, often outcompeting *S. aureus* (3, 9, 12). One factor that contributes to *P. aeruginosa* dominance is the secretion of inhibitory compounds that compromises *S. aureus* growth. These compounds include LasA protease, the siderophore pyoverdine, phenazine toxins, and the quorum-sensing products HQNO and PQS (13–16). However, the competitive dominance of *P. aeruginosa* is not universal especially not in more host relevant contexts. For example, environmental factors such as oxygen levels can influence the dynamics between the two species, with *P. aeruginosa* generally outcompeting *S. aureus* under oxygen-rich conditions, whereas *S. aureus* may persist in anaerobic niches (17). Mutual antagonistic interactions can further be associated with extra benefits, such as increased biofilm formation, which provides enhanced protection from host immune responses and antibiotic treatment (18). Finally, the co-existence of the two species can also occur because the pathogens occupy different niches like in chronic burn wounds infections (4).

The above considerations all refer to ecological interactions without implying any evolutionary change. During repeated interactions between the two species, evolutionary processes matter, and studies revealed that both *S. aureus* and *P. aeruginosa* can adapt to one another (19). Our previous work found that three different *S. aureus* strains quickly adapted and developed resistance to inhibitory *P. aeruginosa* compounds (PQS, HQNO) involved in respiration inhibition and formation of reactive oxygen species (20). However, there were substantial differences between strains. *S. aureus* 6850 showed clear changes in phenotypes – growth restoration, increased staphyloxanthin production, complete resistance to PQS, and higher survival under reactive oxygen species exposure – associated with an accumulation of mutations in transmembrane transporters. In contrast, fewer phenotypic changes and associations with specific mutations were observed in *S. aureus* JE2 although this strain also showed significant growth restoration. This latter finding led us to hypothesize that *S. aureus* JE2 may acquire resistance to inhibitory compounds of *P. aeruginosa* through a combination of phenotypic plastic responses and genetic adaptations. Notably, *S. aureus* JE2 seems particularly suitable to test our hypothesis because it demonstrated higher competitiveness against *P. aeruginosa* PAO1 than other strains (12), suggesting that it has evolved mechanisms to cope with strong competitors.

Here, we tested this hypothesis by combining proteomics, targeted metabolomics and growth assays. Specifically, we compared the responses of an ancestral naive *S. aureus* JE2 strain (henceforth JE2_anc) that had never experienced exposure to *P. aeruginosa* supernatant to the responses of an evolved *S. aureus* JE2 strain (henceforth JE2_evo) that had been exposed to *P. aeruginosa* supernatant for 30 days. For JE2_evo, we took an isolate from our evolution experiment with a mutation in the *tcyA* gene, representing the only mutational change acquired during experimental evolution (20). The *tcyABC* operon encodes an L-cystine importer, and the observed mutation introduces an early stop codon in the gene encoding the substrate-binding domain TcyA. For JE2_anc, we took the ancestral strain from which JE2_evo evolved. We exposed both strains to either tryptic soy broth (TSB) supplemented with spent *P. aeruginosa* PAO1 supernatant or a control condition consisting of TSB only. We compared the proteomes between JE2_anc and JE2_evo to identify evolved differences between strains, whereas a comparison across growth conditions captured phenotypic plastic responses. Subsequently, we allocated differentially regulated proteins to functional classes and used a set of JE2 mutants from a library to assess the benefit of specific regulatory responses.

## Results

### Profound changes in the *S. aureus* proteome in response to *P. aeruginosa* supernatant exposure

To explore the extent to which *S. aureus* alters its proteome when exposed to the spent supernatant of *P. aeruginosa*, we grew the strains *S. aureus* JE2_anc and JE2_evo in 70% tryptic soy broth (TSB) supplemented with 30% spent supernatant of *P. aeruginosa* PAO1 (henceforth called PA supernatant) and compared the proteome to the control condition (growth in 100% TSB). We carried out the experiment in three-fold replication for each condition and *S. aureus* JE2 strain. For our proteome analysis, we included all proteins that we identified based on at least one single matching peptide allowing for a 5% false discovery rate (FDR ≤ 5%, using the target-decoy approach).

This approach successfully identified 1,803 proteins, representing 61.4% of the 2,938 proteins from *S. aureus* JE2. Protein abundances varied profoundly between culturing conditions (*S. aureus* with or without PA supernatant) but only little between strains (JE2_anc vs. JE2_evo) (Fig. 1A). Indeed, a principal component analysis reveals strong clustering of the replicates according to culturing conditions (along PC1, explaining 49.5 % of the total variance, Fig. 1B, Fig. S1). We found 338 (18.7%) proteins to be significantly differentially expressed (q < 0.05, FDR-corrected p-value). Out of these, 335 (99.1%) were differentially expressed between the two culturing conditions, while only 3 proteins were differentially expressed between the two strains and none in response to an interaction between the two factors (Fig. 1C). These results reveal a strong plastic response of *S. aureus* upon exposure to PA supernatant, whereas genetically determined responses acquired during experimental evolution are small.

**Figure 1.**
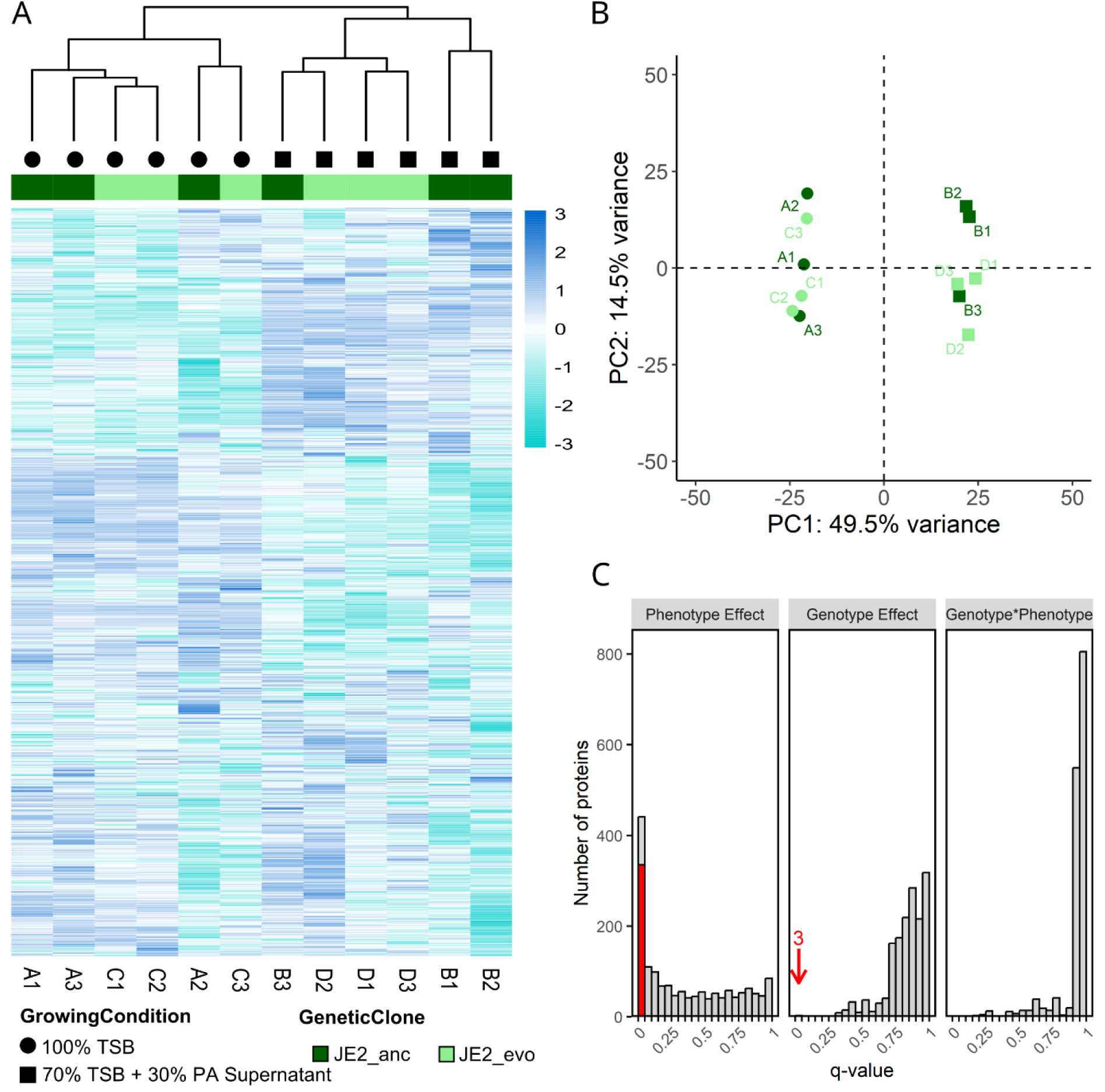
Heatmap and principal component analysis reveal clear proteome differences between culturing conditions (presence/absence of *P. aeruginosa* supernatant) but few differences between strains. **(A)** Heatmap showing hierarchical clustering of protein abundances in JE2_anc (dark green) and JE2_evo (light green) grown either in 70% TSB with 30% PA supernatant (square) or in 100% TSB (circle). Each row represents an individual protein, with the color-code indicating protein abundance (log Z scale), ranging from dark blue (high abundance) to turquois (low abundance). **(B)** Principle component (PC) analysis on the measured protein abundances. The first two PCs account for 64% of the total variation observed. Squares and circles show replicates grown in TSB supplemented with PA supernatant and TSB alone, respectively. Light and dark green symbols depict JE2_evo and JE2_anc replicates, respectively. **(C)** Results of the differential protein expression analysis are shown as distributions of q-values (FDR-corrected p-values) for all proteins, separated by culture condition comparison (Phenotype Effect), strain comparison (Genotype Effect), and their interaction. The red bar and arrow indicate the number of significantly differentially expressed proteins with log2-fold changes > |1|.

### Downregulation of TcyABC transporter proteins as evolutionary response to PA supernatant exposure

We first focus on the proteome difference between JE2_anc and JE2_evo. The latter differs from the former by a single base pair deletion in the *tcyA* gene (position 2’536’157) resulting in an early stop codon due to a frame shift. In accordance with this genetic difference, we found that TcyA and TcyB are the most down-regulated proteins in JE2_evo compared to JE2_anc (Fig. 2A, TcyA: log₂ FC (fold-change) = -5.95, q < 0.0004; TcyB: log₂ FC = - 2.86, q = 0.0012). The genes *tcyA* and *tcyB* are part of an operon (*tcyABC*) together with *tcyC*. We therefore conclude that the stop-codon introducing frame-shift mutation in *tcyA* abolishes the expression of TcyA, down-regulates the expression of TcyB as polar effect, but has a smaller effect on the expression of TcyC (log₂ FC = -1.77, q = 0.3430), the last gene in the operon. In addition to TcyA and TcyB, we identified NTPase as significantly downregulated (Fig. 2A, log₂ FC = -4.45, q = 0.0332). However, in-depth analyses suggest that this hit is a false positive (Fig. S2). We used the deep learning model Koina to predict the spectrum intensities from the identified peptide sequence, and the resulting spectra differed from the actual experimentally measured evidence for this peptide (21).

**Figure 2.**
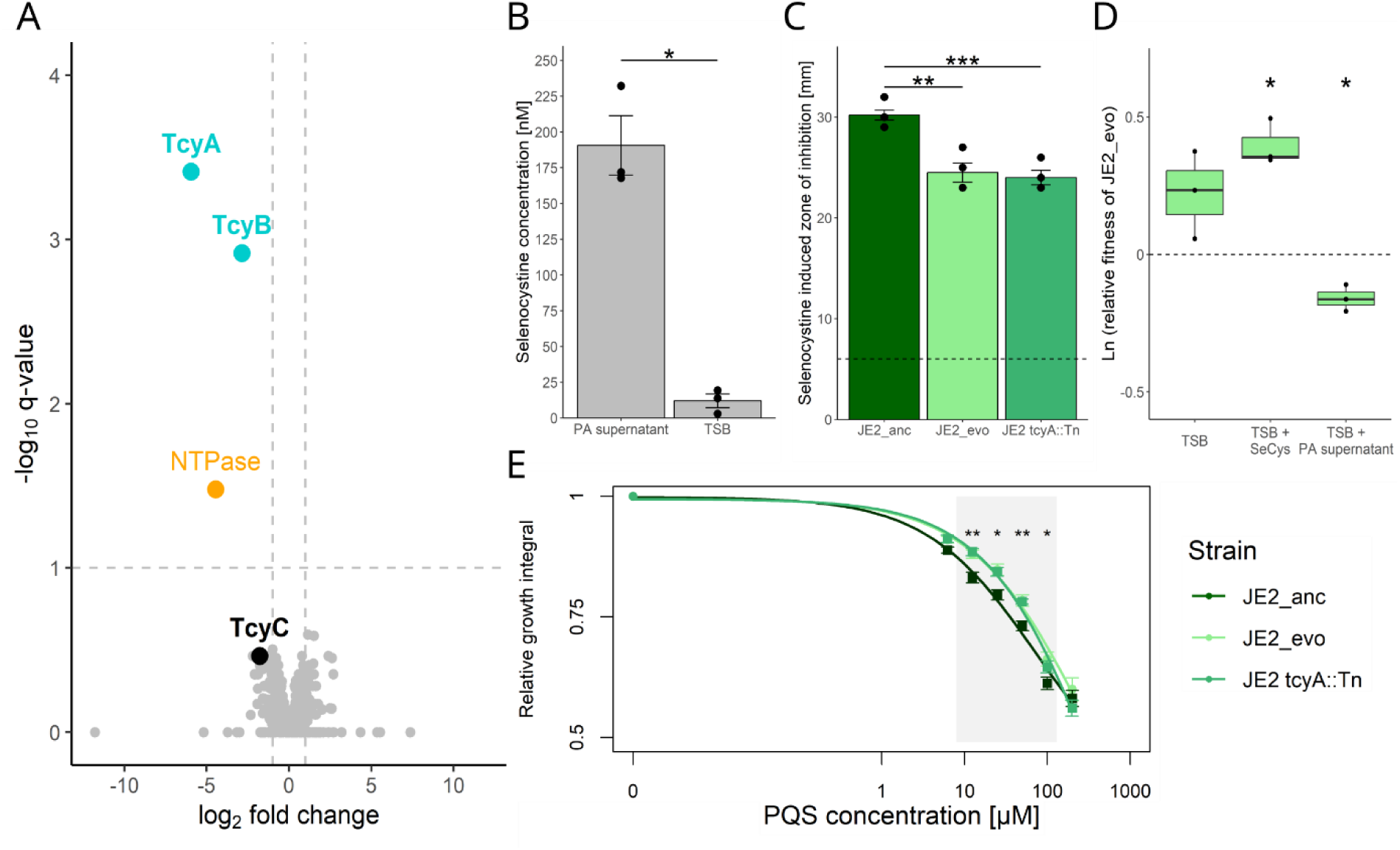
Downregulation of the transporter proteins TcyA and TcyB allow *S. aureus* to grow slightly but significantly better when exposed to selenocystine and PQS produced by *P. aeruginosa*. **(A)** Volcano plot contrasting protein expression level differences (log2) versus q-values (-log10) based on a differential expression analysis between *S. aureus* JE2_anc and JE2_evo (genotype effect). Turquoise dots depict significantly differently regulated proteins (q-values < 0.05 and absolute log2 expression difference >|1|). The orange dot most likely represents a false positive (see main text). **(B)** Targeted metabolomics revealing the presence of significant amounts of selenocystine (SeCys) in PA supernatant and its absence in the TSB medium (Welch’s t-test: t = 8.36, df = 2.21, p = 0.010). **(C)** Zone of inhibition (in mm) for *S. aureus* JE2_anc, JE2_evo and JE2 *tcyA*::Tn when exposed to selenocystine (100 mM) on a paper disk. The dotted line represents the disk diameter (6mm). **(D)** Fitness of JE2_evo relative to JE2_anc after direct competition between the strains in either TSB, TSB with selenocystine or TSB with PA supernatant. Amplicon sequencing was used to assess the frequency of strains after competition. **(E)** Growth dose-response curves of JE2_anc, JE2_evo, and JE2 *tcyA*::Tn when exposed to a gradient of PQS concentrations. Growth trajectories were measured across 10 hours and growth integrals (area under the curve) are expressed relative to growth in the absence of PQS. Asterisks denote significance levels: ***P < 0.001; **P < 0.01; *P < 0.05.

The tcyABC operon encodes a L-cystine transport system, consisting of a substrate-binding domain (TcyA), a transmembrane domain (permease TcyB), and an ATP-binding cassette domain (TcyC). We used the T3PQ approach (22) to estimate the stoichiometry of the tcyABC operon. Our analysis indicates that the *tcyA* gene is more highly expressed in JE2_anc (32-fold increase, Fig. S3) than *tcyB* and *tcyC*. This difference in expression levels hints towards TcyABC being a Type II ATP-Binding Cassette (ABC) transporter, where the substrate binding protein occurs in high excess and does not directly bind to the permease (23). The proteome data of JE2_evo suggests that this strain lacks the substrate-binding domain TcyA, whilst still producing residual amounts of the other two proteins, resulting in a non-functional TcyABC transporter (Fig. S3). Since mutations in *tcyA* repeatedly occurred in response to PA supernatant exposure in our previous experiment (20), we derive that the downregulation of the TcyABC transporter offers a selective advantage to JE2 when exposed to inhibitory compounds secreted by *P. aeruginosa*.

### Non-functional TcyABC transporter confers partial tolerance to selenocystine and PQS

We hypothesized that a non-functional TcyABC transporter could prevent the import of toxic compounds secreted by *P. aeruginosa*. As TcyABC imports cysteine, one possible additional substrate could be selenocystine, a toxic cysteine analogue. *P. aeruginosa* possesses the genes required for selenocysteine production, which forms toxic selenocystine upon oxidation (24–26). We used targeted metabolomics and found that selenocystine is indeed present in the supernatant of PA with an estimated concentration of 190.7 ± 20.8 nM (Figure 2B).

We then used a disk diffusion assay to quantify the susceptibility of JE2 strains to selenocystine. We found that JE2_anc was significantly more inhibited by selenocystine than JE2_evo (Tukey HSD post hoc following one-way ANOVA: mean difference = -5.7, 95% CI = [-8.42, -2.98], p = 0.0005) and a *tcyA* transposon mutant (mean difference = -6.2, 95% CI = [-8.92, -3.48], p = 0.0003; Figure 2C). Important to note is that mutations in *tcyA* only conferred partial resistance to selenocystine and only provided an advantage at high concentrations of this toxic compound. Nonetheless, these results suggest that JE2_evo may experience a selective advantage in direct competition with JE2_anc under selenocystine exposure. To test this, we competed the two strains at equal starting frequencies in either TSB alone, TSB with selenocystine or TSB with PA supernatant. We used amplicon sequencing to assess the frequency of the two strains after 24 hours (Figure 2D). We found that JE2_evo indeed outcompeted JE2_anc in TSB with selenocystine (t_2_ = 8.19, p = 0.0146) but not in TSB alone (t_2_ = 2.41, p = 0.1376). In contrast, JE2_evo had lower relative fitness than JE2_anc in TSB with supernatant (t_2_ = -5.72, p = 0.0293). These results indicate that *tcyA* mutations offer a certain protection against selenocystine resulting in relative growth benefits, but only upon direct selenocystine exposure, and not when exposed to PA supernatant containing a mixture of compounds.

This finding motivated us to explore whether a non-functional TcyABC transporter offers protection against other metabolites secreted by *P. aeruginosa*. We focused on PQS (Pseudomonas Quinolone Signal) because we previously demonstrated that this secreted molecule inhibits naive *S. aureus* strains, while evolved strains showed improved tolerance (20). Here, we exposed JE2_anc, JE2_evo, and JE2::tcyA to a gradient of PQS concentrations over 10 hours (PQS is not stable after 15 hours) (27). We found that JE2_evo and JE2 tcyA::Tn grew significantly better than JE2_anc at intermediate PQS concentrations (12.5 µM – 50 µM), (Figure 2E).

In summary, our results indicate that several toxic compounds secreted by *P. aeruginosa* potentially enter *S. aureus* JE2 via the TcyABC transporter. Mutations leading to a non-functional version of the transporter confer increased tolerance to these toxic compounds.

### *S. aureus* down-regulates virulence factors in response to *P. aeruginosa* supernatant

We now explore the phenotypic plastic proteome changes shown by *S. aureus* in response to PA supernatant exposure (Fig. 1C). Out of the 335 significantly differentially expressed proteins, we found 139 proteins to be upregulated, while 196 proteins were downregulated in TSB with PA supernatant compared to TSB alone (Figure 3A). We were able to map 169 (51%) of the differentially expressed proteins to the KEGG database, while the remaining proteins were either not annotated (42%) or belonged to enzymatic reactions unlinked to specific pathways (7%). When looking at functional pathways, we found that the environmental information processing pathway (proteins, N=142) and the human disease pathway (N=54) were enriched for significantly differentially regulated proteins under PA supernatant exposure (Fig. 3B and Table S1; Fisher’s exact test for, Environmental information processing: odds ratio = 1.66, 95% CI = [1.15, 2.39], p = 0.0461; Human disease: odds ratio = 2.97, 95% CI = [1.72, 5.12], p = 0.0020).

**Figure 3.**
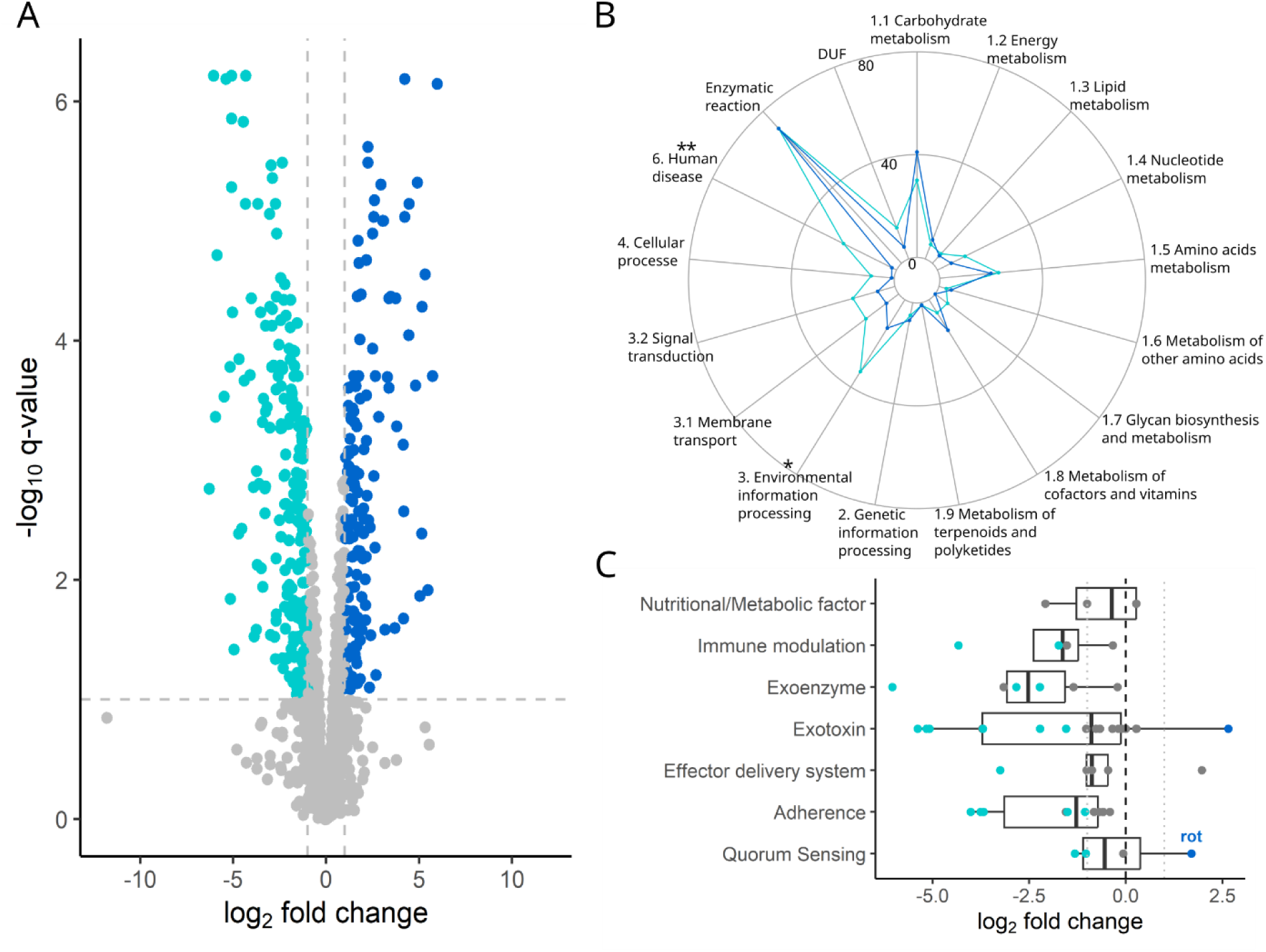
The proteome profile of *S. aureus* changes fundamentally upon exposure to PA supernatant including a major downregulation of virulence traits. **(A)** Volcano plot contrasting protein expression level differences (log2) versus q-values (-log10) based on a differential expression analysis between growth conditions (70% TSB + 30% PA supernatant versus 100% TSB, phenotype effect). Turquoise (n = 196) and dark blue (n = 139) dots depict significantly down- and up-regulated proteins, respectively (q-values < 0.05 and absolute log2 expression difference >|1|). **(B)** Spider chart showing the absolute number of significantly regulated proteins mapped according to the KEGG-pathways (metabolism [split in nine subcategories], genetic information processing, environmental information processing [subcategories: membrane transport, signal transduction], cellular processes, human disease). Further shown are DUF (domains of unknown functions), enzymatic reactions unliked to specific pathways. Asterisks denote pathways enriched for significantly differentially regulated proteins under PA supernatant exposure. **P < 0.01; *P < 0.05. **(C)** Proteins from the human disease pathway (KEGG annotation), grouped according to function based on the virulence factor database (VFDB). Turquoise (n = 20) and dark blue (n = 2) dots depict significantly down- and up-regulated proteins, respectively.

We first turn to the human disease pathway, which includes traits involved with antimicrobial resistance, infection and virulence. Out of the 25 differentially regulated proteins, two were significantly upregulated, while 23 were significantly downregulated in the presence of PA supernatant (Figure 3B). These results clearly indicate that *S. aureus* down-regulates infection and virulence traits in response to inter-species competition. Inspired by these results, we consulted the virulence factor database (VFDB) (28), which features a more specific and complete data set for *S. aureus*. We could map 68 proteins, out of which 30 (44%) were significantly downregulated and two (3%) were significantly upregulated (Figure 3C), reinforcing the view of widespread virulence factor down-regulation. Notably, one of the two upregulated proteins is Rot (repressor of toxins). It plays an important role in *S. aureus* quorum sensing by negatively regulating the expression of genes encoding various exotoxins (29, 30). Among these, we found the LukD (log₂ FC = 5.10, q = 0.0144), LukG (log₂ FC = 5.20, q < 0.0001), and LukH (log₂ FC = 5.40, q < 0.0001) exotoxins (belonging to the Panton-Valentine leukocidins) and the alpha-hemolysin (Hla, log₂ FC = 5.00, q < 0.0001) to be among the most significantly downregulated proteins upon PA supernatant exposure.

In sum, our results reveal an extensive plastic response by *S. aureus* upon exposure to PA supernatant with one of the main responses being the significant downregulation of virulence-related genes.

### *S. aureus* up-regulates ROS scavenging proteins and changes transporter regulation in response to PA supernatant exposure

Next, we examined the environmental information processing pathway, including signal transduction and membrane transport. Since our previous work revealed that HQNO and PQS (promoting the formation of reactive oxygen species, ROS) are involved in *S. aureus* inhibition, we first examined whether *S. aureus* JE2 upregulates ROS scavenging proteins. Indeed, we found the three major proteins SodA (Superoxide Dismutase A, log₂ FC = 2.09, q = 0.0216), KatA (Catalase A, log₂ FC = 1.20, q = 0.0003), and AhpC (Alkyl Hydroperoxide Reductase C, log₂ FC = 2.03, q = 0.0008) to be significantly upregulated. These results demonstrate that *S. aureus* JE2 phenotypically responds to stress induced by ROS.

Another key finding of our previous work was that *S. aureus* strains had many mutations in membrane transporter genes (20). Building on this, we hypothesized that *S. aureus* might exhibit phenotypic plasticity by generally down-regulating transporters when exposed to PA supernatant. We thus mapped our detected proteins to all known *S. aureus* transporters listed in the BioCyc database (31, 32). Out of 282 mapped transporter proteins, eight were significantly upregulated, while 26 were significantly down-regulated (Figure 4A, Table S2) supporting our notion of transporter down-regulation being more common than up-regulation. Among the down-regulated transporters, we find numerous oligopeptide importers and several metal (iron, natrium, nickel) importers (Table S2). Conversely, among the most up-regulated transporters we find sugar(-derivate) importers (gluconate, glucoside, lactose, trehalose), an amino acid importer and EmrB, the transmembrane domain of an efflux pump, known to be involved in *S. aureus* beta-lactam resistance (33). Altogether, these results provide a more nuanced view on *S. aureus* transporter regulation upon PA supernatant exposure, which is characterized by a shift involving down-regulation of oligopeptide and metal importers and upregulating sugar(-derivate) importers and an efflux pump.

**Figure 4.**
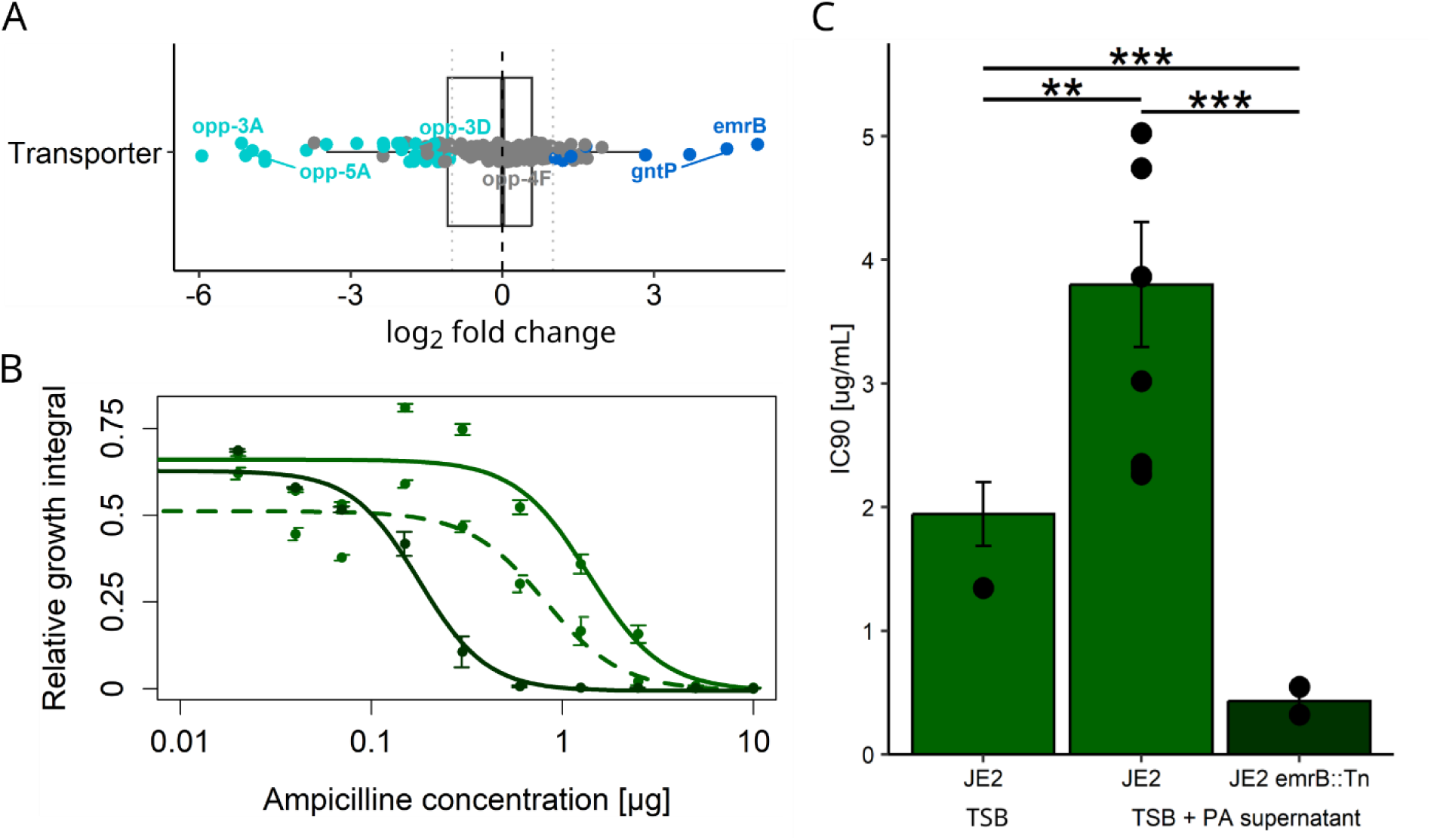
Proteome changes in *S. aureus* upon PA supernatant exposure involve the downregulation of transmembrane transporters and the upregulation of an efflux pump. **(A)** Boxplot showing the differential expression of membrane transporters between growth conditions (70% TSB + 30% PA supernatant versus 100% TSB, phenotype effect). Turquoise (n = 26) and dark blue (n = 8) dots depict significantly down- and up-regulated proteins, respectively (q-values < 0.05 and absolute log2 expression difference >|1|, Table S2). **(B)** Dose-response curves of *S. aureus* strains when subjected to a gradient of ampicillin concentrations (µg/ml). Green circles with dashed line: JE2_anc grown in 100% TSB; green circles with solid line: JE2_anc grown in 70% TSB + 30% PA supernatant; dark green circles with solid line: JE2 *emrB*:Tn grown in 70% TSB + 30% PA supernatant. JE2 *emrB*:Tn is deficient for the EmrB efflux pump, which is up-regulated under PA supernatant exposure and confers increased resistance to toxic compounds. **(C)** Statistical analysis of the IC90-values extracted from the dose-response curves shown in (B). Asterisks denote significant differences: ***P < 0.001; **P < 0.01.

To examine whether these phenotypic responses are beneficial, we focused on EmrB and asked whether the up-regulation of this efflux pump could be involved in expelling toxic compounds from the cell. We used ampicillin as a test probe and predicted that JE2_anc should tolerate higher levels of ampicillin when exposed to PA supernatant compared to when grown in TSB alone because the supernatant increases the expression of the EmrB efflux pump. Our data provide strong support for this hypothesis (Fig. 4B+C). The ampicillin concentration required to inhibit culture growth by 90% (IC90) was significantly higher for JE2_anc when grown in PA supernatant compared to when grown in TSB (fold-change: 1.93, t_19.62_ = -3.27, p = 0.0038). As negative control, we used JE2 emrB::Tn (a transposon mutant deficient for EmrB) and found that this strain was significantly more susceptible to ampicillin compared to JE2_anc in TSB (IC90 fold-change: 4.58, t_8.52_ = -5.52, p = 0.0004). In sum, these results show that the PA-supernatant induced upregulation of the efflux pump EmrB protects *S. aureus* from harmful compounds.

## Discussion

In this study, we investigated the phenotypic responses shown by *S. aureus* JE2 when exposed to the supernatant of its competitor *P. aeruginosa* containing growth inhibitory compounds. Using proteomics, we found that *S. aureus* JE2 significantly down-regulates virulence traits, up-regulates ROS-scavenging systems, changes the expression of transporters (influx is generally reduced, while efflux is increased), and adjusts its nutrient acquisition strategies. Overall, the expression of 18.6% (335) of all identified proteins significantly changed in response to PA supernatant exposure. This profound phenotypic shift strongly contrasted with the minor genetically-induced changes observed in a *S. aureus* JE2 strain (JE2_evo) that had previously been exposed to PA supernatant over 30 consecutive days (20). Compared to its ancestor, we only detected two differentially expressed proteins, TcyA and TcyB, which are both part of a transmembrane importer. Their down-regulation is directly linked to a mutation in *tcyA* acquired during experimental evolution, conferring partial resistance to toxic compounds secreted by *P. aeruginosa*. Altogether, our results indicate that *S. aureus* JE2 has evolved mechanisms to detect threats by competing species and to mount multi-variate plastic responses. These plastic responses are strong and seem to be very effective in protecting *S. aureus* JE2 against toxic compounds secreted by *P. aeruginosa*, limiting the need for genetically hard-wired adaptive strategies arising through mutation and selection.

We propose that the observed proteome changes can be divided into two separate behavioral strategies of how *S. aureus* JE2 can cope with both indirect and direct competition imposed by *P. aeruginosa*. Indirect competition is typically guided by nutrient competition, whereby species do not harm each other directly but compete for the consumption of common resources (34, 35). We interpret the down-regulation of virulence factor production and the shift in nutrient acquisition strategies as measures to cope with nutrient competition. The down-regulation of unneeded virulence factors, such as Panton-Valentine leukocidins that target mammalian leukocytes, would save metabolic costs. A shift in nutrient acquisition strategies, meanwhile, could alleviate nutritional niche competition with *P. aeruginosa* and potentially allow for faster growth.

Contrary to indirect mechanisms, direct competition is typically mediated by secreted or (cell-to-cell) translocated toxic compounds that specifically serve to harm competitors (35). *P. aeruginosa* is particularly potent in secreting toxic compounds, including phenazines (pyocyanin), hydrogen cyanide, rhamnolipids, quinolones (HQNO and PQS), and siderophores (14, 36, 37). Several of these compounds (phenazines, HQNO, PQS) promote the formation of reactive oxygen species (ROS), such that the observed up-regulation of ROS scavengers can be interpreted as a targeted response to the presence of these harmful compounds in the PA supernatant (see also (19)). Furthermore, many of the toxic compounds are secondary metabolites featuring an oligopeptide backbone. Accordingly, the down-regulation of oligopeptide importers (e.g., the Opp-family, Table S2) (38) can be seen as specific responses to limit the uptake of harmful compounds (see (39) for a specific example). Similarly, the up-regulation of the EmrB efflux pump can be seen as responses to expel toxic compounds already taken up. Finally, secreted virulence factors often serve as cues for competitors to launch attacks (40), meaning that the observed down-regulation of virulence factors by *S. aureus* JE2 could reflect a measure to become less detectable by *P. aeruginosa* or elicit a less aggressive response (41). This latter argument is supported by earlier work showing that interactions between *S. aureus* and *P. aeruginosa* is modulated by quorum sensing and secreted virulence factors of both species (11, 42, 43). In sum, our results suggest that *S. aureus* JE2 has evolved fine-tuned mechanisms to sense and react to competitors in a plastic manner.

Our results identify selenocystine as a new potential toxic molecule through which *P. aeruginosa* can inhibit *S. aureus* JE2. *P. aeruginosa* possesses *selD*, a gene encoding an enzyme involved in selenocysteine synthesis (44). Selenocysteine is non-toxic. However, under oxidizing conditions two selenocysteine molecules can form a di-selenide bond resulting in the synthesis of toxic selenocystine (45). The formation of selenocystine is promoted by ROS and we found that selenocystine indeed accumulates in the PA supernatant in significant amounts. These biochemical processes help to better understand the phenotypic and genetic responses of *S. aureus* JE2 to PA supernatant. At the phenotypic level, the significant upregulation of ROS-scavenging proteins could be both a general response to competition and a specific response to reduce the formation of selenocystine. Whereas the superoxide dismutase A (SodA) catalyzes the conversion of superoxide anion (O₂⁻) into hydrogen peroxide (H₂O₂) and oxygen (O₂), the catalase KatA and the peroxiredoxin AhpC convert hydrogen peroxide to water, oxygen, or alcohols (46–48). At the genetic level, the selection of *tcyA* mutations in experimentally evolved *S. aureus* JE2 lines (20) suggest a second layer of defense by reducing the uptake of selenocysteine or selenocystine into the cell. However, *tcyA* mutations may also incur a cost as cysteine uptake (important for sulphur metabolism) is compromised (26). Relevant in this context is the role of cysteine as building block for the synthesis of thiol-containing molecules such as glutathione, molecules that are involved in ROS deactivation (49–51). The putative costs associated with *tcyA* mutations could explain why JE2_evo only outcompeted JE2_anc when exposed to selenocystine alone but not when exposed to PA supernatant containing compounds promoting ROS formation.

Our analysis reveals that plasticity in protein expression can be an adaptive trait itself (52). In our case, plasticity allows *S. aureus* JE2 to sense the presence of a competitor and mount protective measures to guarantee survival and growth. The benefit of such plastic responses is that they can be switched off when not needed, whereas costs can accrue through the maintenance of sophisticated sensing mechanisms and taking erroneous decisions, i.e. mounting a response when not needed. Clearly, plastic responses differ from genetically hard-wired responses such as the permanent loss of the TcyABC-transporter in our JE2_evo strain. Whether or not natural selection favors plastic versus hard-wired genetic adaptations probably depends on ecological parameters, such as the rate of environmental fluctuation. Genetic adaptation may be favored under strong and permanent selection pressures, for example under antibiotic exposure. Conversely, phenotypic plasticity may be favored under weaker and fluctuating selection. The latter scenario certainly applies to interspecies interactions, where episodes of mono- and mixed-species growth cycles may alternate.

In summary, we observed a major change in the proteome of *S. aureus* JE2 when this pathogen was exposed to the supernatant of its competitor *P. aeruginosa*. Most changes can intuitively be interpreted as phenotypically plastic adaptations to the threat imposed by the typically dominant *P. aeruginosa*. Plastic responses involve the downregulation of virulence factors and transmembrane importers, the upregulation of an efflux pump and ROS scavenging proteins, and a shift in the expression of nutrient importers. These strong plastic response raises two major questions. First, can *S. aureus* JE2 build up memory such that the level of protection increases upon repeated exposure to PA supernatant? Such memory effects could explain why *S. aureus* JE2 cultures were no longer inhibited in PA supernatant after 30 days of exposure even in the absence of genetic beneficial mutations. Second, do plastic phenotypic changes proceed genetic adaptation? This concept is a matter of constant debate in evolutionary biology (53–56) with increasing support accumulating in various organisms including microbes (57–61). Taken together, we show that plastic changes in protein expression are a major component guiding inter-species interactions. In future studies, we need to obtain a deeper understanding of how such plastic effects affect species co-existence and virulence in acute and chronic polymicrobial infections (11).

## Material and Methods

### Bacterial strains and culture conditions

We used four *S. aureus* JE2 strains: JE2_anc is our wild type strain, while JE2_evo is a clone from our previous experimental evolution study (clone-ID: B0604), in which JE2 was exposed to PA supernatant for 30 consecutive days (20). During its evolution, JE2_evo acquired a single point mutation in the *tcyA* gene (position 2’536’157) resulting in an early stop codon due to a frame shift. Otherwise, JE2_anc and JE2_evo are isogenic. We further used the transposon mutants JE2 *tcyA*::Tn and JE2 *emrB*::Tn, both obtained from the Nebraska transposon mutant library (Table S3; (62)). We further utilized *P. aeruginosa* PAO1 (ATCC 15692) to generate supernatant.

We stored all strains in tryptic soy broth (TSB) containing 40% glycerol at -80°C degrees. All growth experiments were conducted in TSB medium (Becton Dickinson, Heidelberg, Germany). We grew overnight cultures in 4 mL TSB within 13 mL tubes at 37°C and 170 rpm. The next day, we washed overnight cultures with 0.8% sodium chloride (NaCl) solution and adjusted them to an optical density at 600 nm (OD600) of 0.4. For our growth assays, we inoculated these cells into 96-well plates containing the corresponding medium, to obtain a starting OD600 of 0.0004. For all growth assays (including supernatant assays, PQS and ampicillin-inhibition assays), we incubated the 96-well plates at 37°C in a plate reader (Tecan Infinite M Nano, Tecan, Männedorf, Switzerland) and measured strain growth by recording OD600 every 5 minutes for 24 hours, with 30 seconds shaking events prior and after to recordings.

### Sample preparation for proteome analysis

We generated cell-free supernatants by growing *P. aeruginosa* overnight in 10 mL of TSB within 50 mL Falcon tubes. The cultures were incubated at 37°C with aeration at 220 rpm. The following day, we centrifuged the cultures and filtered-sterilized the supernatants using 0.2 µM pore size filters (Whatman, Fisher Scientific, Reinach, Switzerland). In parallel, we grew JE2_anc and JE2_evo in TSB for 18 hours. Subsequently, we centrifuged and washed the bacterial cell with 10 mL 0.8% NaCl. We then subjected the cells of the two strains (1 mL, adjusted to a final OD600 to 0.004) to two growth conditions (9 mL) in three-fold replication each: 100% TSB as the regular medium or 70% TSB mixed with 30% PA supernatant. After 18h of growth at 37°C and 170 rpm, we centrifuged the cultures, washed and resuspended the pellets with 0.8% NaCl.

### Protein extraction and protein digestion

From the above-generated samples, we each used 100 µl to obtain pelleted cells through centrifugation at 3000 g for 2 min. After supernatant removal, 200 µl of lysis buffer (4 % Sodium dodecyl sulfate (SDS) in 100 mM Tris/HCl pH 8.2) were added per sample. Protein extraction was carried out using a tissue homogenizer (TissueLyser II, QUIAGEN) by applying 2×2min cycles at 30 Hz. The samples were treated with High Intensity Focused Ultrasound (HIFU) for 1 minute at an ultrasonic amplitude of 90 % before boiling at 95°C for 10 minutes while shaking at 800 rpm on a Thermoshaker (Eppendorf). After sample cool down 10 U of Benzonase (Sigma-Aldrich Chemie GmbH) were added per sample and incubated for 45 min at 37°C. Protein concentration was determined using the Lunatic UV/Vis polychromatic spectrophotometer (Unchained Labs) with a 1:50 dilution for each sample. For each sample 100 µg according to Lunatic measurement were taken and reduced with 5 mM TCEP(tris(2-carboxyethyl)phosphine) and alkylated with 15 mM chloroacetamide at 60°C for 30 min.

Samples were processed using the single-pot solid-phase enhanced sample preparation (SP3). The SP3 protein purification, digest and peptide clean-up were performed using a KingFisher Flex System (Thermo Fisher Scientific) and Carboxylate-Modified Magnetic Particles (GE Life Sciences; GE65152105050250, GE45152105050250) (63, 64). Beads were conditioned following the manufacturer’s instructions, consisting of 3 washes with water at a concentration of 1 µg/µl. Samples were diluted with 100% ethanol to a final concentration of 60% ethanol. The beads, wash solutions and samples were loaded into 96 deep well- or micro-plates and transferred to the KingFisher. Following steps were carried out on the robot: collection of beads from the last wash, protein binding to beads, washing of beads in wash solutions 1-3 (80% ethanol), protein digestion (overnight at 37°C with a trypsin:protein ratio of 1:50 in 50 mM Triethylammoniumbicarbonat (TEAB)) and peptide elution from the magnetic beads using MilliQ water. The digest solution and water elution were combined and dried to completeness and re-solubilized in 20 µL of MS sample buffer (3% acetonitrile, 0.1% formic acid). The samples were diluted 1:5 with MS sample buffer spiked with iRT peptides (Biognosys AG).

### Liquid chromatography-mass spectrometry analysis

Mass spectrometry analysis was performed on an Orbitrap Fusion Lumos (Thermo Scientific) equipped with a Digital PicoView source (New Objective) and coupled to a M-Class UPLC (Waters). Solvent composition of the two channels was 0.1% formic acid for channel A and 0.1% formic acid, 99.9% acetonitrile for channel B. For each sample 1 µl was loaded on a commercial MZ Symmetry C18 Trap Column (100Å, 5 µm, 180 µm x 20 mm, Waters) followed by nanoEase MZ C18 HSS T3 Column (100Å, 1.8 µm, 75 µm x 250 mm, Waters). The peptides were eluted at a flow rate of 300 nl/min. After an initial hold at 5% B for 3 min, a gradient from 5 to 22% B in 80 min and 32% B in 10 min was applied. The column was washed with 95% B for 10 min and afterwards the column was re-equilibrated to starting conditions for an additional 10 min. Samples were acquired in a randomized order. The mass spectrometer was operated in data-dependent mode (DDA) acquiring a full-scan MS spectra (300−1’500 m/z) at a resolution of 120’000 at 200 m/z after accumulation to a target value of 500’000. Data-dependent MS/MS were recorded in the linear ion trap using quadrupole isolation with a window of 0.8 Da and HCD fragmentation with 35% fragmentation energy. The ion trap was operated in rapid scan mode with a target value of 10’000 and a maximum injection time of 50 ms. Only precursors with an intensity above 5’000 were selected for MS/MS and the maximum cycle time was set to 3 s. Charge state screening was enabled. Singly, unassigned, and charge states higher than seven were rejected. Precursor masses previously selected for MS/MS measurement were excluded from further selection for 20 s, and the exclusion window was set at 10 ppm. The samples were acquired using internal lock mass calibration on m/z 371.1012 and 445.1200. The mass spectrometry proteomics data were handled using the local laboratory information management system (LIMS) (65, 66). All relevant data have been deposited to the ProteomeXchange Consortium via the PRIDE (http://www.ebi.ac.uk/pride) partner repository with the data set identifier PXD059647.

### Protein identification, label free protein quantification and statistical analysis

The acquired raw MS data were processed by MaxQuant (version 1.6.2.3), followed by protein identification using the integrated Andromeda search engine (67). Spectra were searched against the subsequently mentioned strain specific SA JE2 protein database, concatenated to its reversed decoyed fasta database to estimate false discovery rates. Carbamidomethylation of cysteine was set as fixed modification, while methionine oxidation and N-terminal protein acetylation were set as variable. Enzyme specificity was set to trypsin/P allowing a minimal peptide length of 7 amino acids and a maximum of two missed-cleavages. MaxQuant Orbitrap default search settings were used. The maximum false discovery rate (FDR) was set to 0.01 for peptides and 0.05 for proteins. Label-free quantification was enabled and a 2-minutes window for the match between runs was applied. In the MaxQuant experimental design template, each file is kept separate in the experimental design to obtain individual quantitative values. The proteinGroups.txt file was used as input to statistical analysis and differential expression.

The R package prolfqua (68) was used to normalize and further analyze the differential expression among conditions. The proteinGroups.txt file generated by MaxQuant was used directly. In brief, the protein intensities were first normalized for each sample with a modified robust z-score transformation. In prolfqua two factors were defined as “growing condition,“ either TSB or PASN, and “genotype,” either ancestor or evolved clone. A linear model was fitted for each protein to explain the normalized protein abundance using a combination of the two factors. Different contrasts were calculated for each protein to compare the growing condition effect or the genotype effect. In the case of complete missing values in one condition, the group average was estimated using the mean of 10% of the smallest protein intensities.

To estimate the stoichiometry of the TcyABC transporter, we initially analyzed the peptides.txt file generated by MaxQuant, ensuring accurate peptide-to-protein roll-up. Subsequently, we used the R package prolfqua for further analysis. For data aggregation, we utilized the mean_topN method, selecting N=3 to focus on the three most intense peptides per protein. This approach allowed us to retrieve reliable quantitative protein values based solely on the most significant peptide data.

### Generation of strain-specific protein databases

A JE2 strain specific protein sequences were translated from the ncbi genome from GenBank (https://www.ncbi.nlm.nih.gov/nuccore/CP020619.1/) using the ncbi service by sending the coding sequences as protein format to a file. These were 2938 protein sequences. Since the identifier type that is used in this fasta file is unknown to most tools available for downstream analysis, a multiblast with this JE2 strain specific fasta file was performed against the uniprot reference proteome (https://www.uniprot.org/proteomes/UP000008816), downloaded in June 2021) and the description lines of the JE2 strain specific sequences were extended with the orthologue identifier of the best blastp match of the *S. aureus* uniprot reference proteome along with the corresponding blastp e-value. Furthermore, some protein sequences annotated in the corresponding gff file as pseudogenes or frame shifted genes were also translated in all 6 frames in the annotated regions as specified in the gff file. This resulted in additional 720 protein sequences where stop codons were replaced with X (any amino acid). Also 495 protein sequences of known contaminants were included in this database. In total our generated protein database consists of 4154 protein sequences.

### Proteomics functional categorization

We mapped and categorized all detected proteins to their corresponding Kyoto Encyclopedia of Genes and Genomes (KEGG) pathways (69, 70). These pathways are grouped according to their biological functions. For our analysis, we included five main pathways Metabolism, Genetic Information Processing, Environmental Information Processing, Cellular Processes, and Human Disease, and subdivisions of them to gain an in-depth understanding of the exact functions of the proteins.

### Targeted Metabolomics

To quantify selenocystine concentrations in either the TSB control or the *P. aeruginosa* supernatant we added 400µl of ice cold 100% (v/v) MeOH to 100µl of our samples. After vortexing briefly and incubation for 10min on ice the samples were centrifuged (13.2krpm/+4°C/10min). 450µl of each supernatant were transferred to new 1.5ml Eppendorf PP tubes and dried under a gentle stream of nitrogen. The dried residues were reconstituted in 100µl H_2_O containing 0.1% (v/v) formic acid and 0.05% (v/V) heptafluorobutyric acid and submitted to LC-MS analysis.

Selenocystine was quantified by Selected Reaction Monitoring (SRM) based reversed phase capLC-nanoESI-MSMS analysis on a triple-quadrupole type mass spectrometer (TSQ Quantiva, Thermo Fisher Scientific Inc., CA, USA). Acquired SRM data were analyzed by using the QuanBrowser module of the Xcalibur software (Thermo Fisher Scientific Inc., CA, USA). For each sample, absolute quantities of the metabolite was reported, based on the co-analysis of a dilution series of the pure compound.

### Disk diffusion assays

To measure the potency of selenocystine to inhibit the various JE2 strains, we followed the methodology described by (26). Briefly, we spread an overnight culture of either JE2_anc, JE2_evo, or JE2 tcyA::Tn on petri dishes containing 20 ml of tryptic soy agar (TSA) to allow the growth of an extensive bacterial lawn. Subsequently, we prepared sterile Whatman paper discs soaked with 10 µl selenocystine solutions at concentrations of 100mM, 50mM, or 10mM. These solutions were prepared in 1N HCl. The discs were then placed on TSA dishes (five-fold replication), and incubated at 37°C for a period of 24 hours. The bacterial growth inhibition was measured by the diameter of the clear zone surrounding the disc, recorded in millimeters.

### Competition assay

We used amplicon sequencing to distinguish between JE2_anc and JE2_evo in our competition assays, as the two strains only differ by a single nucleotide deletion in the *tcyA* gene. Competition between the two strains was carried out in three different media (200 µl volumes) with three-fold replication each: 100% TSB, 70% TSB mixed with 30% spent PA supernatant and 100% TSB containing 200 nM selenocystine. This concentration was chosen to match the experimentally measured concentration of selenocystine in PA supernatants. We used a two-step PCR approach and performed a first step PCR after 24h using primers targeting the *tcyA* gene (Table S4). The second PCR step attached sequencing adapters and sample-specific barcodes (indices) to the amplicons generated in the first PCR. This step enables sample identification and compatibility with the Illumina sequencing platform, allowing for efficient multiplexing in a single run. The amplified DNA was then sent for commercial library preparation and sequencing. Sequencing was conducted on the Illumina MiSeq Platform, with 2×250 bp paired-end reads targeting 200,000 reads per sample. For 7 samples, we obtained 100,000 to 110,000 reads, and for 2 samples, only a few thousand reads were obtained (these were samples with supernatant). Independent of the number of reads achieved, we observed similar relative fitness across samples. Raw sequencing data were demultiplexed and trimmed of Illumina adaptor residuals, and the absence/presence of the nucleotide deletion was used to quantify the relative abundance of JE2_anc and JE2_evo.

### PQS assays

To test whether JE2_anc, JE2_evo, and JE2 tcyA::Tn differ in their susceptibility to PQS, we exposed the three strains to a gradient of PQS concentrations and measured their growth. Specifically, we filled a round-bottom 96-well plate with 180 µL of TSB. We then prepared the following working concentrations of PQS (heptyl-3-hydroxy-4(1H)-quinolone, Sigma-Aldrich, Buchs, Switzerland) in DMSO: 2000 µM, 1000 µM, 500 µM, 250 µM, and 125 µM. From these working stocks, we added 10 µL to the corresponding wells. As negative control, we added 10 µL of DMSO to the medium. Finally, we added 10 µL of bacterial culture (prepared as described above) of one of the three strains to each well. We conducted the experiment independently five times, incubating the 96-well plates at 37°C degrees in a plate reader (Tecan Infinite M Nano, Tecan, Männedorf, Switzerland) and measured strain growth by recording optical density at 600nm (OD600) every 5 minutes for 24 hours, with 30 seconds shaking prior and after each recording.

### Determination of ampicillin dose-response curves

To test whether the upregulation of the EmrB efflux system protects JE2 from toxic compounds, we subjected JE2 strains to a range of ampicillin concentrations. We used the broth micro dilution method with two different media: a mix of 40% TSB + 60% NaCl (promoting normal EmrB expression) and a mix of 40% TSB + 60% spent PA supernatant (promoting increased EmrB expression). The broth micro dilution assay is carried out in 96-well plates with each well containing 100μL medium. To the first column, we added 100μL of ampicillin solution (final concentration: 20 μg/ml). We then serially diluted the medium-antibiotic mix by passing 100μL from the first column to the subsequent nine columns resulting in an antibiotic concentration gradient covering 10μg, 5μg, 2.5μg, 1.25μg, 0.625μg, 0.313μg, 0.156μg, 0.078μg, 0.039μg, 0.020μg of ampicillin per ml. No antibiotics were added to the eleventh column. Next, we prepared bacterial cultures using the strains JE2_anc and JE2 *emrB*::Tn (as described above). We then added 10μL of culture to all wells, except those in the twelfth column, which served as blank media controls (10 μL of 0.8% saline solution was added instead). Experiments featured three replicates and were repeated three times on different days. For all experiments, we measured OD600 every 5 minutes over 24 hours at 37°C in a Tecan Infinite M Nano plate reader, with 30 seconds of shaking before and after each recording.

### Statistics

All statistical analyses were performed in R 4.1.2 and RStudio version 2024.09.1+394. A protein is considered significantly differently regulated if its Log2 fold change is either less than -1 or greater than 1, combined with a false discovery rate FDR ≤ 0.05. These criteria ensure that the observed changes in protein abundance changes are both statistically significant and biologically meaningful. Whenever necessary, we applied the Benjamini-Hochberg correction to adjust for multiple comparisons, using the false discovery rate (FDR) method.

We used a two-sample t-test to determine whether selenocystine is detectable and present in the PA supernatant but not in the control medium. We applied a one-way ANOVA with Tukey’s HSD post hoc test to compare whether the radius of the inhibition zones caused by selenocystine in the disk diffusion assay differs across JE2 strains. To check whether the ln-transformed relative fitness of JE2_evo (in comparison to JE2_anc) significantly differs from zero (the expected value for fitness parity), we used one-sample t-tests. To compare whether growth under different PQS concentrations differ across JE2 strains, we used Welchs’s ANOVA (to account for the inequality of the variances). To establish dose-response curves for ampicillin, we fitted four-parameter logistic functions to the growth data (measured as area under the curve of growth trajectories). From these fits, we extracted the IC90-value (the compound concentration required to inhibit bacterial cultures by 90%) for each strain and growth medium and then compared the IC90-values using multiple t-tests.

## Data availability statement

All proteomic data have been deposited to the ProteomeXchange Consortium via the PRIDE (http://www.ebi.ac.uk/pride) partner repository with the data set identifier PXD059647. All other data supporting the findings of this study are available from the corresponding author upon request.

## Acknowledgements

This project has received funding from the Swiss National Science Foundation (grants no 31003A_182499 and 310030_212266) to R.K., and from the University of Zurich Teaching Fund (2019_12/Fostering OMICS research through research-based teaching and learning) to J.G. and L.P. Conceptualization was done by the following: L.S., S.N., R.K. and J.G. Data curation was done by the following: L.S., R.K. and J.G. Formal analysis was done by the following: L.S., N.Z., W.W., C.P., J.G. Investigation was done by the following: L.S., S.N., J.G., and W.W. Methodology was done by the following: L.S., S.N., N.Z., W.W., C.P. and J.G. Visualization was done by the following: L.S. J.G. Writing— original draft was done by the following: L.S.. R.K. and J.G. Writing—review and editing was done by the following: L.S., S.N., N.Z., W.W., C.P., R.S., R.K. and J.G. All authors read and approved the final manuscript.

