## Supplemental Material for "A combination of phenotypic responses and genetic adaptations enables *Staphylococcus aureus* to withstand inhibitory molecules secreted by *Pseudomonas aeruginosa*"

This document contains the following supplementary material:

- 3 supplemental Figures
- 4 supplemental Tables

**Supplemental Figure S1**

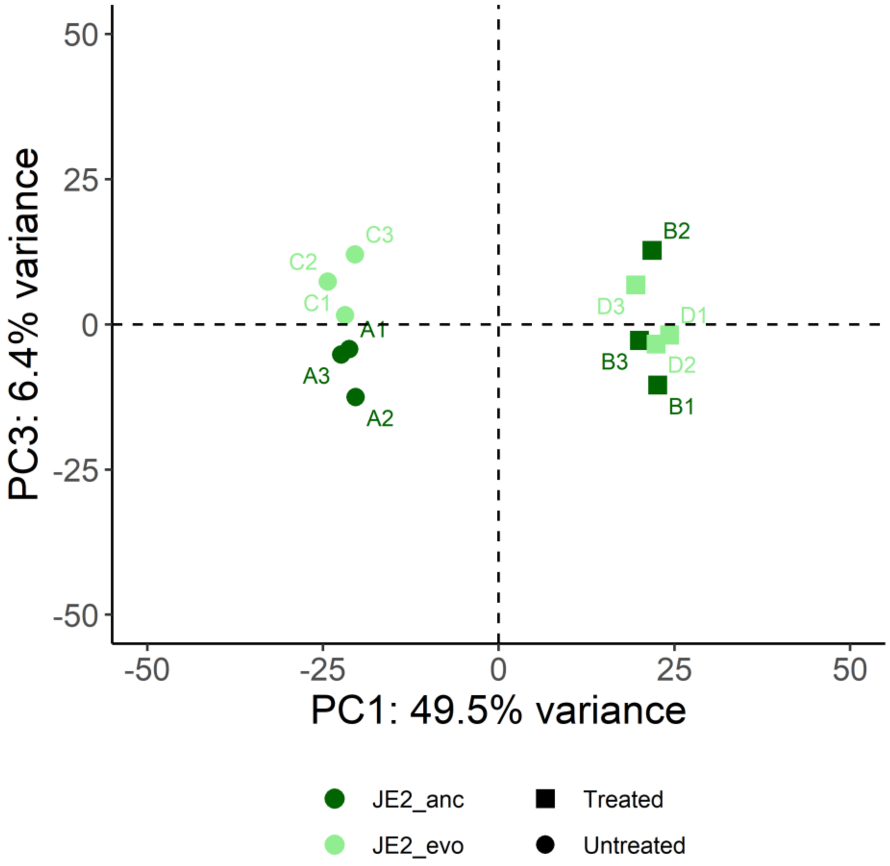


**Figure S1.** Principle component (PC) analysis on all measured proteins showing the first and third PC. They account together for 55.9% variation observed in the data. Squares and circles differentiate replicates grown in either 70% tryptic soy broth (TSB) supplemented with 30% spent supernatant of *P. aeruginosa* PAO1 (square) or 100% TSB medium (circle). Light green symbols: JE2_evo. Dark green symbols: JE2_anc.

**Supplemental Figure S2**

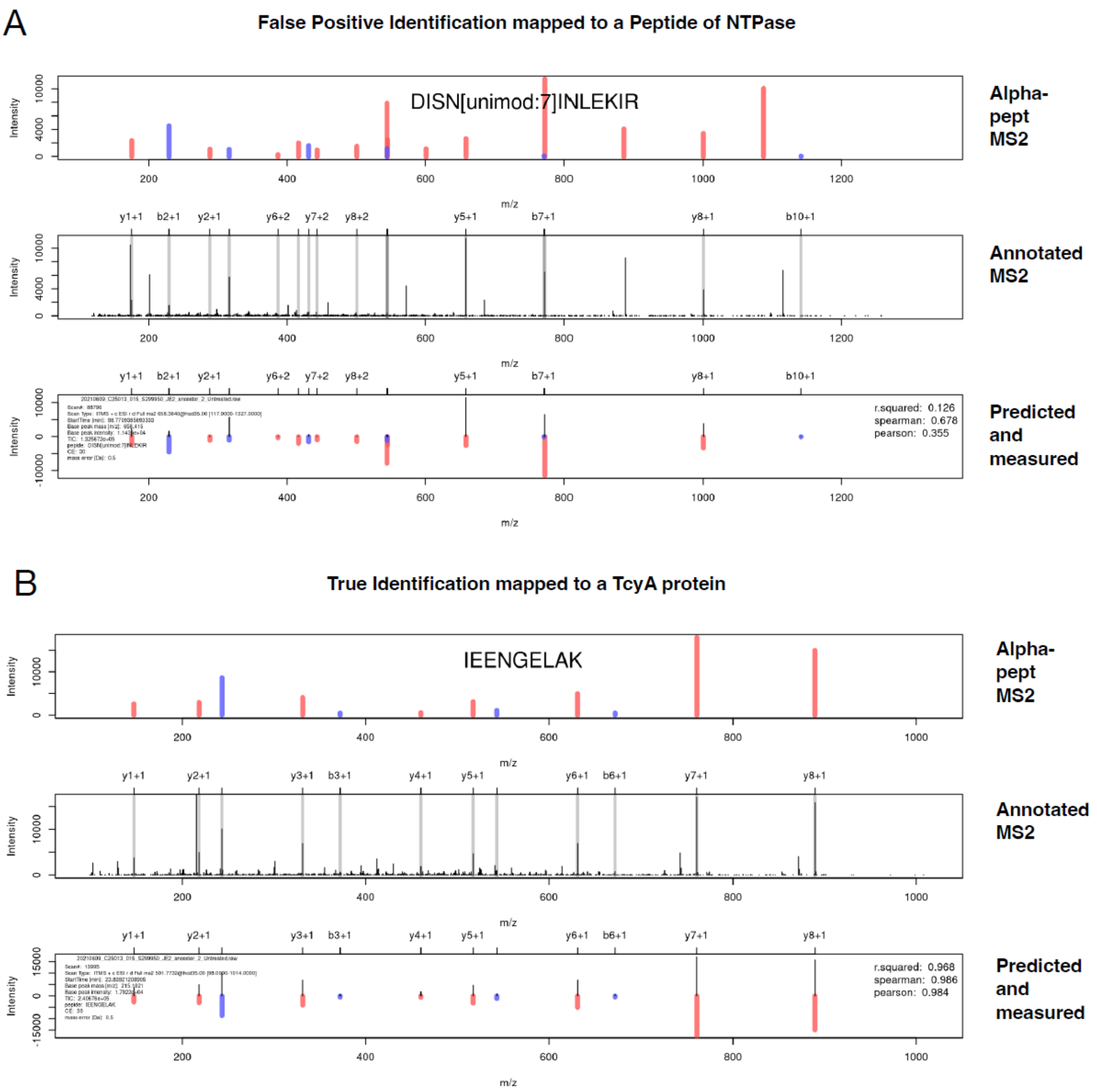


**Figure S2.** To illustrate the false positive match of the DISN[unimod:7] INLEKIR peptide, we predicted the spectrum using the AlphaPept_ms2_generic model, which is available through the Koina model repository. As an initial step, we conducted a collision energy (CE) normalization using true positive (TP) hits, such as the peptide IEE shown in panel B. The resulting CE value of 30 eV was then used as input together with the peptide sequence and the charge state to predict the DISNI peptide spectrum (Gessulat et al., 2019; Zeng et al., 2022). Panel A displays the evidence for the identification of the NTPase protein, which was determined to be a false positive. Panel B shows a valid identification for a peptide from the TcyA protein. In both panels, the top section illustrates the predicted y-ion (red) and b-ion (blue) spectra generated by koina (Alpha-pept MS2). The middle section presents the experimentally measured and annotated spectrum with matched y- and b-ions (Annotated MS2). The bottom section features a mirror plot comparing the experimental and predicted spectra for the matched ions (Predicted and Measured).This allows to calculate correlation statistics to identify true from false positive identifications.

**Supplemental Figure S3**

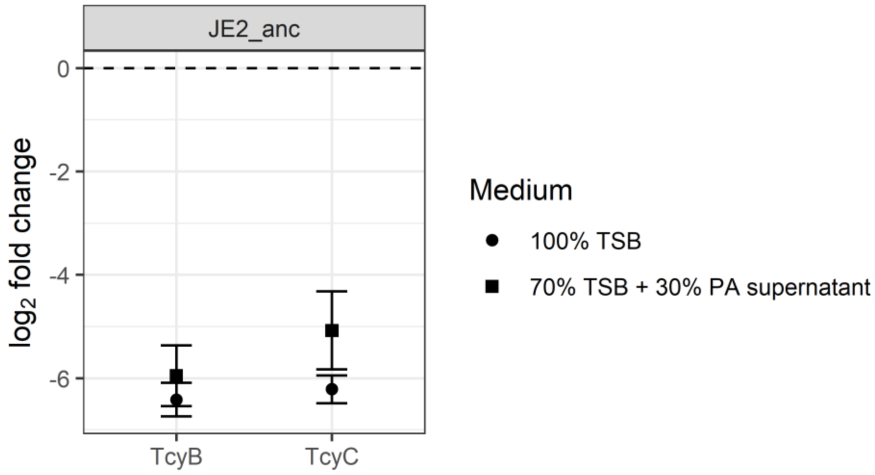


**Figure S3.** Log-fold intensity change of TcyB and TcyC relative to TcyA for JE2_anc, grown in either 100% TSB or 70% TSB + 30% PA supernatant. The analysis shows that TcyA is much more abundant than TcyB and TcyC, indicating that TcyABC is a Type II ATP-Binding Cassette transporter, where the substrate binding protein TcyA occurs in high excess and does not directly bind to the permease TcyB.

**Supplemental Table S1** KEGG Pathways

| Pathway | Total Proteins | Non-Significant regulated | Significant upregulated | Significant downregulated | BH-adjusted p-value |
|---|---|---|---|---|---|
| 6. Human Disease | 54 | 29 | 2 | 23 | 0.0020 |
| 3. Environmental Information Processing | 142 | 96 | 13 | 33 | 0.0461 |
| 3.2 Signal transduction | 72 | 48 | 7 | 17 | 0.1600 |
| 4. Cellular Processes | 26 | 16 | 1 | 9 | 0.2200 |
| 3.1 Membrane transport | 70 | 48 | 6 | 16 | 0.2200 |
| 1.1 Carbohydrate metabolism | 272 | 201 | 41 | 30 | 0.3310 |
| 1.5 Amino acid metabolism | 163 | 121 | 20 | 23 | 0.4440 |
| 1.6 Metabolism of other amino acids | 31 | 23 | 5 | 3 | 0.8390 |
| 1.2 Energy metabolism | 57 | 43 | 8 | 6 | 0.8390 |
| 1.8 Metabolism of cofactors and vitamins | 94 | 74 | 14 | 6 | 1 |
| 1.7 Glycan biosynthesis and metabolism | 31 | 25 | 0 | 6 | 1 |
| DUF | 91 | 73 | 5 | 13 | 1 |
| NA | 765 | 599 | 66 | 100 | 1 |
| 1.3 Lipid metabolism | 57 | 48 | 4 | 5 | 1 |
| 1.4 Nucleotide metabolism | 104 | 86 | 6 | 12 | 1 |
| 1.9 Metabolism of terpenoids and polyketides | 28 | 26 | 1 | 1 | 1 |
| 2. Genetic Information Processing | 171 | 159 | 7 | 5 | 1 |
| Enzymatic Reaction | | | 12 | 12 | Not a KEGG pathway |

**Supplemental Table S2** Transporter genes differently expressed in *S. aureus* grown in PAO1 supernatant vs TSB

| Gene | Fold change (log2)[a] | BioCyc Annotation | Primary Function |
|---|---|---|---|
| emrB | 5.04 | MFS transporter | Exporter |
| gntP | 4.43 | Gluconate permease | Importer |
| brnQ1 | 3.70 | Branched-chain amino acid transporter | Importer |
| glvC | | PTS alpha-glucoside transporter subunit | Importer |
| | 2.83 | IIBC | Importer |
| Q2G144_STAA8 | 1.65 | PTS lactose transporter subunit IIB | |
| treP | 1.35 | PTS trehalose transporter subunit IIBC | Importer |
| htsA | 1.19 | Heme ABC transporter substrate-binding protein | Importer |
| mntA | 1.04 | Phosphonate ABC transporter ATP-binding protein | Importer |
| cntA | -1.05 | Nickel ABC transporter | Importer |
| Q2FVB4_STAA8 | -1.05 | Lantibiotic ABC transporter ATP-binding protein | Importer |
| sirA | -1.27 | Iron ABC transporter substrate-binding protein | Importer |
| rnd2 | -1.31 | Multidrug transporter | Exporter |
| potD | -1.36 | Spermidine ABC transporter substrate-binding protein | Importer |
| metN | -1.51 | ABC transporter ATP-binding protein | Importer |
| Q2FVG0_STAA8 | -1.54 | ABC transporter ATP-binding protein | Importer |
| murP | -1.72 | Permease | Importer |
| dltD | -1.73 | D-alanyl-lipoteichoic acid biosynthesis protein | Not categorized |
| Q2FYU8_STAA8 | -1.76 | Gamma-aminobutyrate permease | Importer |
| opuBB | -1.83 | Glycine/betaine ABC transporter permease | Importer |
| nsaA | -1.98 | ABC transporter ATP-binding protein | Exporter |
| opp3F | -1.99 | Oligopeptide ABC transporter ATP-binding protein | Importer |
| yvrC | -2.02 | ABC transporter substrate-binding protein | Importer |
| opp3D | -2.08 | ABC transporter ATP-binding protein | Importer |
| gmpC | -2.35 | ABC transporter substrate-binding protein | Importer |
| nptA | -2.36 | Na/Pi cotransporter | Importer |
| mscL | -2.89 | Mechanosensitive ion channel protein MscL | Not categorized |
| phnD | -3.49 | Phosphonate ABC transporter substrate-binding protein | Importer |
| crtQ | -3.88 | 4,4-diaponeurosporenoate glycosyltransferase | Not categorized |
| Q2G1F5_STAA8 | -4.70 | Predicted ABC transporter of a peptide | Importer |
| dacA | -4.71 | TIGR00159 family protein | Not categorized |
| Q2FZM4_STAA8 | -4.95 | ABC transporter substrate-binding protein | Importer |

| | | | |
|---|---|---|---|
| Opp4A | -5.08 | Nickel ABC transporter substrate-binding protein | Importer |
| opp3A | -5.16 | Peptide ABC transporter substrate-binding protein | Importer |
| epiF | -5.95 | Lantibiotic ABC transporter ATP-binding protein | Exporter |
| SAUPAN001056000 potential orthologue oppF | MS/MS evidence that protein is only expressed in untreated condition. Peptide intensity is weak but can consistently only be found in untreated conditions | | Importer |

[a] Fold change values are the average of 3 replicates after normalization

**Supplemental Table S3** Strains used for this study.

| Species and strain name | Origin | Description | Reference |
|---|---|---|---|
| *Pseudomonas aeruginosa* (PA) | | | |
| PAO1 | Wound | Commonly used *Pseudomonas aeruginosa* laboratory strain. | ATCC 15692 |
| *Staphylococcus aureus* (SA) | | | |
| JE2_anc | Skin and soft tissue infection | USA300 CA-MRSA isolate. Highly virulent, cytotoxic, and hemolytic. | NARSA |
| JE2_evo | Skin and soft tissue infection | USA300 CA-MRSA isolate. Highly virulent, cytotoxic, and hemolytic, Framshift mutation in the *tcyABC* operon | Niggli et al. 2023 (B0604) |
| JE2 ***tcyA***::Tn | Skin and soft tissue infection | USA300 CA-MRSA isolate. Highly virulent, cytotoxic, and hemolytic, Tn insertion in the *tcyA* gene | Nebraska transposon mutant library |
| JE2 ***emrB***::Tn | Skin and soft tissue infection | USA300 CA-MRSA isolate. Highly virulent, cytotoxic, and hemolytic, Tn insertion in the *emrB* gene | Nebraska transposon mutant library |

CA-MRSA: Community-acquired methicillin-resistant *S. aureus*

**Supplemental Table S4** Primers used for amplicon sequencing

| Target Gene | Encoded Protein | Sequence | Amplicon (bp) |
|---|---|---|---|
| tcyA | Amino acid ABC transporter substrate-binding protein | F:5-GCATGTTGTTGTTCGCTATT-3 | 333 |
| | | R:5-GCTAAGGATAAAGGTGCTGA-3 | |
